# Endogenous TDP-43 prevents retrotransposons activation through Dicer-2 activity and the RNA silencing machinery in Drosophila neurons

**DOI:** 10.1101/604421

**Authors:** Giulia Romano, Raffaella Klima, Fabian Feiguin

## Abstract

The aberrant expression of retrotransposable elements (RTEs) was observed in different neurodegenerative diseases including amyotrophic lateral sclerosis (ALS), a terminal disorder characterized by functional alterations in the small RNA-binding protein TDP-43, suggesting that these events might be connected. Using genome wide gene expression profiles, we detected RTEs highly upregulated in TDP-43-null Drosophila heads while, the genetic rescue of TDP-43 function reverted these modifications. Furthermore, we found that TDP-43 modulates the small interfering RNA (siRNA) silencing machinery responsible for RTEs repression. Molecularly, we observed that TDP-43 regulates the expression levels of Dicer-2 by direct protein-mRNA interactions *in vivo*. Accordingly, the genetic or pharmacological recovery of Dicer-2 activity was sufficient to repress retrotransposons activation and revert the neurodegeneration in TDP-43-null Drosophila motoneurons. Our results, describe a novel physiological role of endogenous TDP-43 in the prevention of RTEs-induced neurodegeneration through the modulation of Dicer-2 activity and the siRNA pathway.

## INTRODUCTION

Amyotrophic lateral sclerosis (ALS) is a devastating disease that affects the homeostasis of the motor system, defined by motoneurons and the associated glia, leading to muscles denervation, progressive paralysis and neurodegeneration. Regarding the pathological mechanisms of the disease, studies performed in brain tissues obtained from deceased patients revealed the presence of insoluble aggregates of the small ribonuclear protein TDP-43 distributed along the cytoplasm and outside the cell nucleus (Arai et al., 2006; Geser et al., 2009; Neumann et al., 2006). These modifications, strongly correlate with the symptoms of the disease and were observed in the great majority of the sporadic and familial cases of ALS (Sreedharan et al., 2008). However, is still a matter of debate how histological alterations in TDP-43 lead to neurodegeneration. In this direction, experiments performed in transgenic animals indicated that TDP-43 is an aggregation prone protein that induce neurodegeneration when overexpressed in neuronal tissues (Cannon et al., 2012; Igaz et al., 2011; Shan et al., 2010; Tsai et al., 2010; Wils et al., 2010; Xu et al., 2010). Moreover, analogous research lines showed that TDP-43 variants carrying mutations linked to familial cases of ALS were more predisposed to form aggregates and, in addition, more neurotoxic (Janssens et al., 2013; Stallings et al., 2010; Swarup et al., 2011; Tian et al., 2011; Wegorzewska et al., 2009; Xu et al., 2011). On the other hand, the formation of insoluble aggregates may also disrupt the physiological function of the endogenous protein and lead to neurodegeneration through mechanisms related with the absence of TDP-43 function in the nucleus. In relationship with these observations, we demonstrated that the suppression of the TDP-43 homolog protein in Drosophila (TBPH), faithfully reproduced in flies the main characteristics of the human disease alike paralysis, motoneurons degeneration and reduced life span (Feiguin et al., 2009; Godena et al., 2011). Moreover, we described that TBPH function is permanently required in neurons and glia to maintain the molecular organization of the neuromuscular synapses as well as prevent the denervation of the skeletal muscles (Romano et al., 2015, 2014), supporting the idea that deficiencies in TBPH function may conduct to ALS by interfering with the physiological regulation of critical metabolic pathways inside the motor system. In order to identify these molecules, we performed a transcriptome comparison of gene expression profiles between wildtype and TBPH null mutant adult head tissues. Intriguingly, we observed that the absence of TBPH provoked the upregulation of notorious families of conserved retrotransposons that included the endogenous retrovirus (ERV) *gypsy.* In addition, we found that the genetic recovery of TBPH activity prevented the activation of these elements, revealing that the endogenous function of TBPH is required for retrotransposons repression. In the present study, we tested the hypotheses described above and explored the mechanisms regulated by TBPH in retrotransposons silencing. Moreover, we investigated the neurological consequences of ERV activation in TBPH-null flies and examined if similar regulatory pathways are conserved in human neuroblastoma cells. Finally, we tested novel pharmacological compounds and therapeutic strategies to compensate the defects of TBPH loss of function in the repression of retrotransposons activation. We hope that our results will provide novel arguments to understand the disease process and facilitate the way to novel curative interventions in ALS.

## RESULTS

### The lack of TBPH induce the expression of retrotransposons in Drosophila

We have previously indicated that the molecular function of TBPH is permanently required in Drosophila motoneurons to prevent muscles denervation, locomotive defects and early neurodegeneration (Romano et al., 2014). In order to identify the molecules involved in the neurodegenerative process initiated by the absence of TBPH, we utilized Drosophila *melanogaster* to analyze differences in the patterns of gene expression between wildtype and TBPH minus flies. For these experiments, the mRNAs expressed in adult heads of TBPH-null alleles (tbph^Δ23^ and tbph^Δ142^) and wildtype controls were isolated to hybridize GeneChip Drosophila Genome 2.0 Arrays. Intriguingly, the statistical analysis of these experiments revealed that 12 out of the 79 transposons, present in the microarray, appeared dysregulated in TBPH-minus alleles compared to wildtype (Figure 1A and Figure 1-figure supplement 1). Interestingly, we observed that the great majority of these transposable elements belonged to the long terminal repeat (LTR) family of retrotransposons. In particular, we found that *accord* and *gypsy* were the most upregulated LTRs in TBPH mutants (Figure 1A). The modifications described in the microarray, were independently confirmed by quantitative RT-PCR (qRT-PCR) using different combinations of primers against the RNA sequences transcribed from these elements (Figure 1B). In addition we observed that the glycoprotein *env*, codified by *gypsy* (Song et al., 1994; Teysset et al., 1998; Touret et al., 2014), emerged upregulated in TBPH-minus heads compared to controls demonstrating by a different methodology that the activity of this retrotransposon was increased in mutant tissues (Figure 1C). More importantly, we found that the genetic expression of the TBPH protein was able to repress the activation of *accord* and *gypsy* in TBPH mutant backgrounds, demonstrating that the role of TBPH was rather specific (Figure 1B and C).

**Figure 1.**
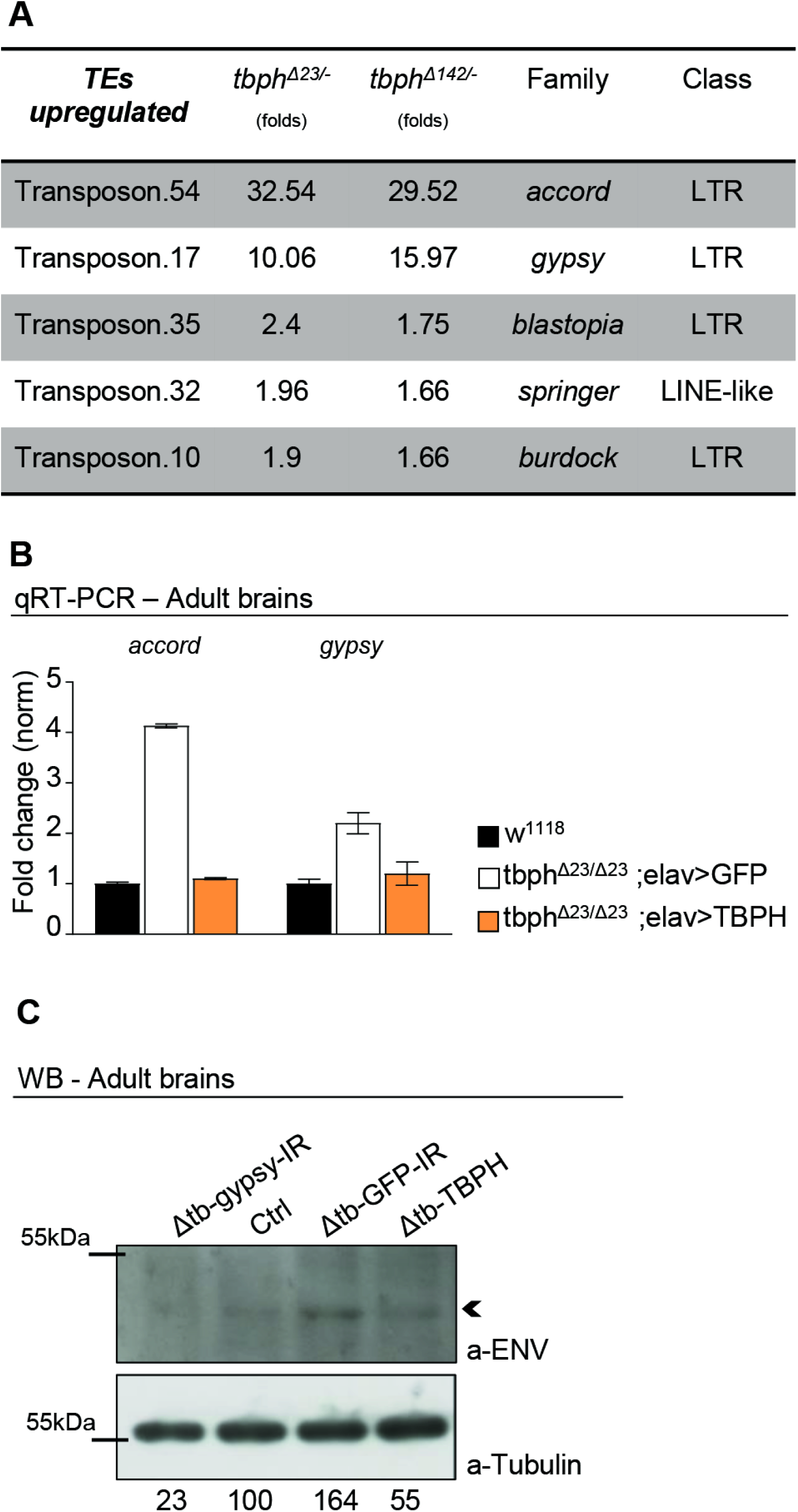
RTEs are upregulated in TBPH mutants. (**A**) Microarray results showing upregulated TEs in TBPH-null mutants: the fold changes are reported for both tbph mutant alleles (Δ23 and Δ142) and referred to *w*^1118^; TEs family and class were also indicated. *n*=3 (biological replicates). (**B**) Real time quantitative PCR reveals *accord* and *gypsy* transcript levels normalized on *Rpl11* (housekeeping) in *w*^1118^ - tbph^Δ23^,elav-GAL4/tbph^Δ23^; UAS-GFP/+ and tbph^Δ23^,elav-GAL4/tbph^Δ23^,UAS-TBPH. *n*=3 (biological replicates, with 3 technical replicates for each), error bars SEM. (**C**) Western blot analysis of Drosophila *env* levels in Δtb-gypsy-IR (tbph^Δ23^,elav-GAL4/tbph^Δ23^; UAS-gypsy-IR/+), ctrl (tbph^Δ23^,elav-GAL4/ +), Δtb-GFP-IR (tbph^Δ23^,elav-GAL4/tbph^Δ23^; UAS-GFP-IR/+) and Δtb-TBPH (tbph^Δ23^,elav-GAL4/tbph^Δ23^,UAS-TBPH). Adult brains, 1 day old, were probed with anti-ENV and alpha-tubulin antibodies. The same membrane was probed with the two antibodies and the bands of interest were cropped. Quantification of normalized amounts was reported below each lane. *n*=2 (biological replicates).

### The activation of retrotransposons causes motoneurons degeneration in TBPH-null flies

The observations related above indicate that the endogenous function of the TBPH protein must be required to prevent the activation of retrotransposons *in vivo*. Furthermore, the data suggests that the mobilization of these elements may contribute to the phenotypes induced by the absence of TBPH activity in Drosophila neurons. To test these possibilities, we treated TBPH-null flies with different combinations of nucleoside and non-nucleoside revert transcriptase inhibitors (NRTI and NNRTI) (Usach et al., 2013). These compounds, are antiretroviral inhibitors that prevent the replication of endogenous retrotransposons by interfering with the enzymatic activity of the reverse-transcriptase or, behaving as chain terminators. As a result, we noticed that the oral administration of the NRTIs: stavudine, azidotimidine, tenofovir and abacavir, together with the NNRTI rilpivirine were able to revert the locomotive defects described in TBPH-minus flies during larvae development (Figure 2A-B and Figure 2-figure supplement 1A). In addition, we decided to analyze more in details the neurological consequences of *gypsy* upregulation in TBPH-minus Drosophila. This is because *gypsy* is a very active retrotransposon in Drosophila, responsible for the majority of the spontaneous mutations described in flies (Krug et al., 2017; Li et al., 2013; Misseri et al., 2004; Song et al., 1994) and, moreover, *gypsy* presents strong similarities with the viral protein HERV-K, a human endogenous retrovirus recently detected in patients with ALS (Douville and Nath, 2017; Li et al., 2015, 2012). Therefore, to test the role of *gypsy* in TBPH-null phenotypes we decided to silence the expression of this retrotransposon in tbph^Δ23^ homozygous flies. For these experiments, we utilized transgenic flies carrying RNAi constructs against the endogenous mRNA sequence of *gypsy* (*gypsy*-IR) cloned in UAS expression vectors (31). Consequently, we found that the neuronal expression of two independent RNAi lines against *gypsy* (*gypsy*-IR_3_ and IR_4_), utilizing the pan-neuronal driver *elav-GAL4* or the more restricted motoneuronal promoter *D42-GAL4*, were able to significantly revert the locomotive phenotypes observed in TBPH-minus third instar larvae (tbph^Δ23^/tbph^Δ23^; *elav-GAL4* or *D42-GAL4*/*gypsy*-IR_3_-IR_4_) compared to analogous flies expressing an RNAi against GFP (tbph^Δ23^/tbph^Δ23^; *elav-GAL4* or *D42-GAL4*/GFP-IR) (Figure 2C). Surprisingly, we noticed that the genetic rescue of the locomotive behaviors induced by the suppression of *gypsy* in TBPH-null backgrounds was followed by the regrowth of the presynaptic terminals and the recovery of the glutamate receptors clusters present at the postsynaptic membranes (Figure 2D-G), demonstrating that the abnormal activation of *gypsy* negatively contributes to the maintenance of the neuromuscular synapses and muscles innervation. Subsequently, we noticed that the suppression of *gypsy* in glial cells, using *repo-GAL4* (tbph^Δ23^/tbph^Δ23^; *repo-GAL4*/*gypsy*-RI_3_), was not able to modify the degenerative phenotypes provoked by the lack of TBPH (Figure 2-figure supplement 1B) suggesting that *gypsy* may not be active in these tissues or, alternatively, the suppression of *gypsy* expression in the glia was not sufficient to prevent the neurodegeneration.

**Figure 2.**
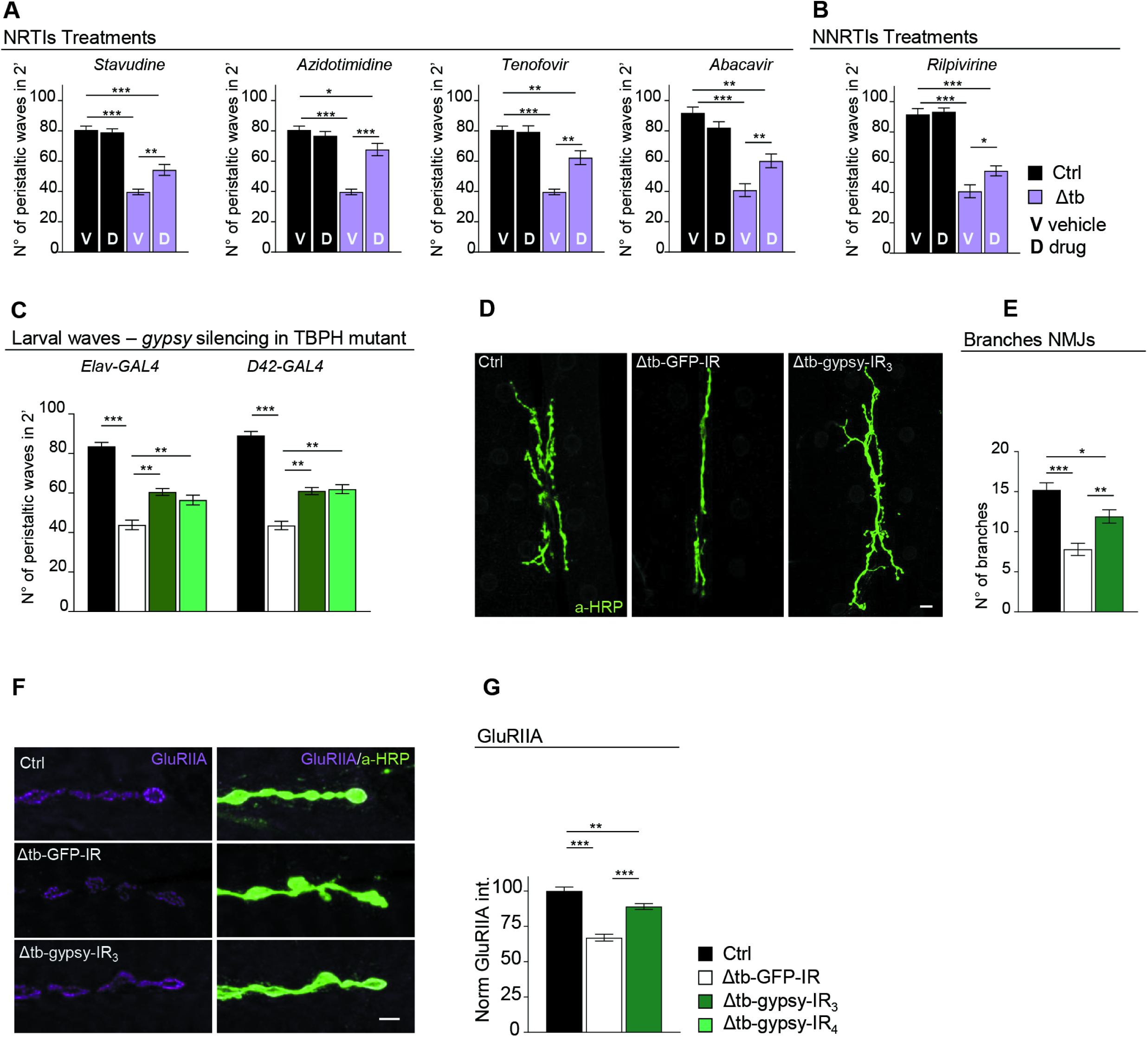
Pharmacological and genetic suppression of RTEs revert TBPH mutant phenotypes. (**A**) Number of peristaltic waves of Ctrl (*w*^1118^) and Δtb (tbph^Δ23^/ tbph^Δ23^) fed with NRTIs drugs (D) compared to vehicle only (V). *n*=20. (**B**) Number of peristaltic waves of Ctrl (*w*^1118^) and Δtb (tbph^Δ23^/ tbph^Δ23^) fed with NNRTIs drugs (D) compared to vehicle only (V). *n*=20. (**C**) Number of peristaltic waves of Ctrl (*w*^1118^), Δtb-GFP-IR (tbph^Δ23^,elav-GAL4/tbph^Δ23^; UAS-GFP-IR/+), Δtb-gypsy-IR_3_ (tbph^Δ23^,elav-GAL4/tbph^Δ23^; UAS-gypsy-IR_3_/+) and Δtb-gypsy-IR_4_ (tbph^Δ23^,elav-GAL4/tbph^Δ23^; UAS-gypsy-IR_4_/+). *n*=20. (**D**) Confocal images of third instar NMJ terminals in muscle 6/7 second segment stained with anti-HRP (in green) in Ctrl, Δtb-GFP-IR and Δtb-gypsy-IR_3_. (**E**) Quantification of branches number. *n*=15. (**F**) Confocal images of third instar NMJ terminals in muscle 6/7 second segment stained with anti-HRP (in green) and anti-GluRIIA (in magenta) in Ctrl, Δtb-GFP-IR and Δtb-gypsy-IR_3_. (**G**) Quantification of GluRIIA intensity. *n*>200 boutons. *p<0.05, **p<0.01, ***p<0.001 calculated by one-way ANOVA, error bars SEM. Scale bar 10µm (in D) and 5µm (in F).

### TBPH controls retrotransposons silencing by regulating Dicer-2 levels

The retrotransposons, including *gypsy*, have the capacity to transcribe themselves through RNA intermediates (Ito and Kakutani, 2014; McCullers and Steiniger, 2017). In physiological conditions, the expression of these elements is maintained under repression by the synthesis of small interference RNAs, in charged to mediate the post-transcriptional silencing of the retrotransposons through the formation of RNA-induced silencing complexes (RISC) (Slotkin and Martienssen, 2007). These siRNAs, present a typical size of 21-23 nucleotides and complementary sequences against different retrotransposons were found to be conserved in different species, as well as, present in different somatic tissues including the brain (Carthew and Sontheimer, 2009; Tabach et al., 2013). Taking in consideration that the expression levels of *gypsy* were upregulated in TBPH-null flies, we decided to test whether the siRNA silencing machinery was affected by the lack of TBPH compared to wildtype controls. For these experiments, we took advantage of a previously described methodology based on the co-expression of a GFP-IR construct together with a GFP reporter in transgenic flies (Krug et al., 2017; Tang et al., 2004).

Thus, differences in the expression levels of the GFP reporter were quantified by western blot and reflected the efficiency of the RNA silencing machineries in different tissues and genetic backgrounds. Accordingly, we utilized *D42-GAL4* to express the constructs described above and observed that wildtype neurons were able to silence more efficiently the GFP reporter compared to TBPH-minus brains, suggesting that the absence of TBPH may have affected the normal functioning of the siRNA machinery (Figure 3A). In order to identify the molecular mechanisms behind these alterations, we investigated whether the expression levels of the different components of the siRNA machinery were affected by the absence of TBPH in Drosophila mutant heads. For these experiments, we utilized qRT-PCR technics to test the brain levels of the principal constituents of the RISC complex like: Dicer-2, loquacious and Argonaute 2. In addition, we analyzed the expression amounts of a different group of genes previously associated with LTR silencing such as piwi, pasha, and homeless (Kavi et al., 2005). Interestingly, our study found that the mRNA levels of RNase Dicer-2 (*Dcr-2*) where the only transcript significantly downregulated in two independent loss of function alleles of TBPH (Figure 3B and Figure 3-figure supplement 1). Furthermore, we observed that the protein levels of *Dcr-2* were similarly downregulated in TBPH-minus heads compared to controls (Figure 3C). In addition, the presence of putative binding sites for TBPH in the coding sequence of *Dcr-2*, decided us to explore whether these molecules physically interact *in vivo*. For these experiments, we expressed a flag-tagged isoform of TBPH in Drosophila neurons and performed pull down assays from fly heads tissues (Godena et al., 2011; Romano et al., 2014). In this manner, we found that the mRNA of *Dcr-2* appeared highly enriched in TBPH immunoprecipitated samples compared to similar experiments performed utilizing a modified variant of this protein that is unable to bind the RNA (TBPH^F/L^), (Ayala et al., 2005; Buratti and Baralle, 2001), confirming that these molecules physically relate *in vivo* (Figure 3D). Additionally, we observed that TBPH was also capable to bind *Dcr-2* at the protein level demonstrating direct protein-protein interactions between these molecules in Drosophila neurons (Figure 3E). Interestingly, we found that the suppression of TDP-43 in human neuroblastoma SH-S5Y5 cells produced a similar reduction in the expression levels of the human protein Dicer suggesting that these regulatory mechanisms must be conserved among the species (Figure 3F), (Kawahara and Mieda-Sato, 2012). Altogether, our results stipulate that TBPH may regulate the proficiency of the siRNA machinery in Drosophila neurons by modulating the expression levels of *Dcr-2* or, alternatively, TBPH could control *Dcr-2* activity through the formation of RISC complexes using direct physical interactions and conserved mechanisms.

**Figure 3.**
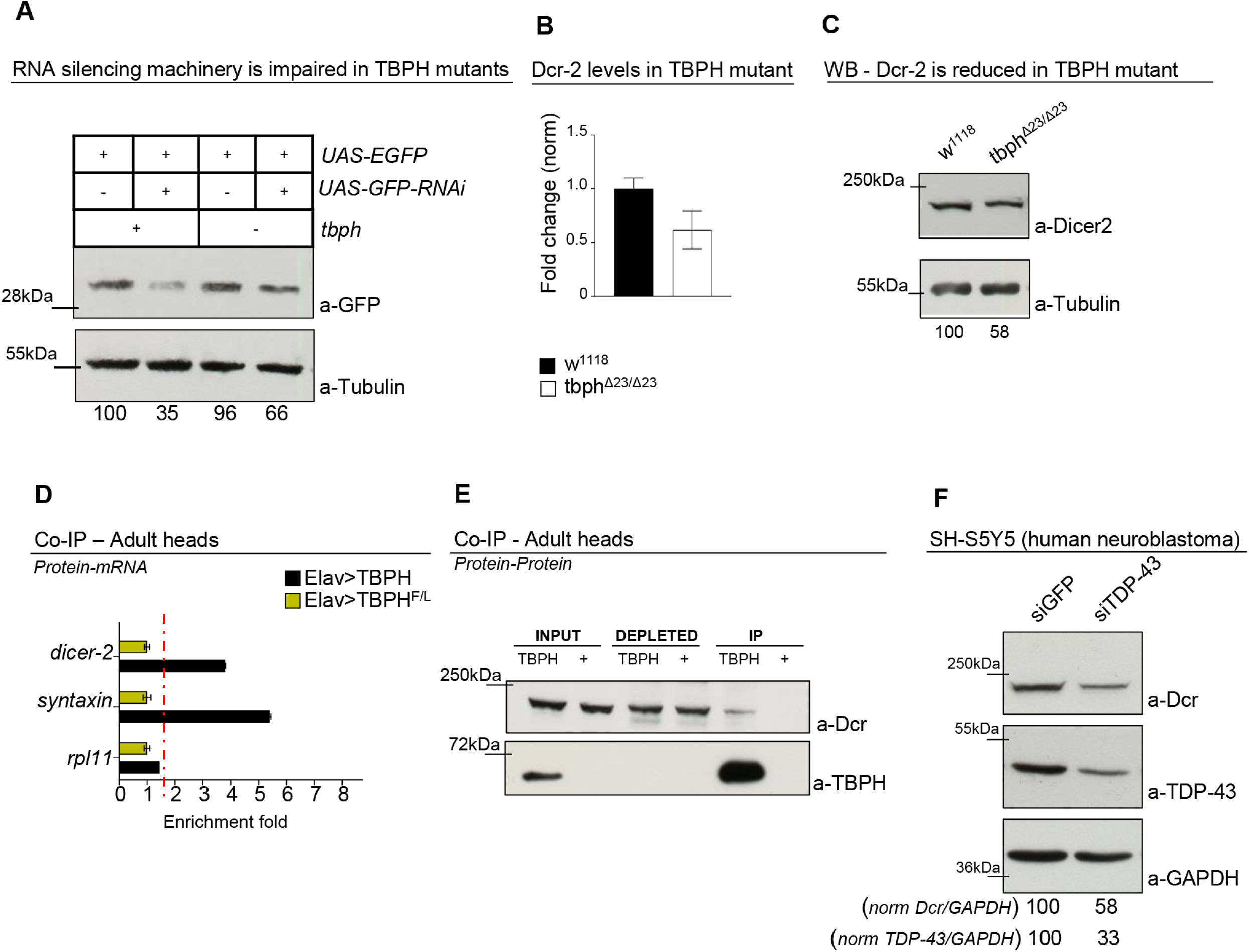
TBPH physically interacts and influences *Dcr-2* levels. (**A**) Western blot analysis of lane 1 (+/+;D42-GAL4,UAS-EGFP/+), lane 2 (+/+;D42-GAL4,UAS-EGFP/UAS-GFP-IR), lane 3 (tbph^Δ23^/tbph^Δ23^; D42-GAL4,UAS-EGFP/+) and lane 4 (tbph^Δ23^/tbph^Δ23^; D42-GAL4,UAS-EGFP/UAS-GFP-IR). Adult brains, 1 day old, were probed with anti-GFP and alpha-tubulin antibodies. The same membrane was probe with the two antibodies and the bands of interest were cropped. Quantification of normalized amounts was reported below each lane. *n*=2 (biological replicates). (**B**) Real time PCR of *Dcr-2* transcript levels normalized on *Rpl11* (housekeeping) in adult heads of *w*^1118^ and tbph^Δ23^/tbph^Δ23^. *n*=3 (biological replicates, with 2 technical replicates for each), error bars SEM. (**C**) Western blot analysis of *w*^1118^ and tbph^Δ23^/tbph^Δ23^ adult brains probed with anti-Dicer and alpha-tubulin antibodies. The same membrane was probe with the two antibodies and the bands of interest were cropped. Quantification of normalized amounts was reported below each lane. *n*=3 (biological replicates). (**D**) qRT-PCR analysis of mRNAs immunoprecipitated by Flag-tagged TBPH (Elav>TBPH) and its mutant variants TBPH ^F/L^ (Elav> TBPH ^F/L^). The *dicer-2* enrichment-folds was referred to *rpl-11* (negative control), syntaxin has been used as positive control. *n*=3 (biological replicates). (**E**) Western blot analysis of proteins immunoprecipitated by Flag-tagged TBPH in adult heads of TBPH (GMR-GAL4/UAS-TBPH) and + (GMR-GAL4/+). Input, depleted and immunoprecipitated materials (IP) were analyzed, probing the membrane with anti-TBPH and Dicer. *n*=2 (biological replicates). (**F**) Western blot analysis on human neuroblastoma (SH-S5Y5) cell line probed for anti-Dicer, anti-GAPDH and anti-TDP-43 in siGFP (GFP ctrl) and siTDP-43 (TDP-43 silenced). The same membrane was probe with the three antibodies and the bands of interest were cropped. Quantification of normalized protein amount was reported below each lane, *n*=3 (biological replicates).

### The rescue of *Dcr-2* expression levels retrieves retrotransposons silencing and recuperates motoneurons degeneration in TBPH-minus flies

The data described above, indicates that endogenous TBPH is physiologically required to prevent neurodegeneration by blocking the activation of retrotransposons through the regulation of *Dcr-2* activity in Drosophila neurons. In order to test this hypothesis, we decided to reestablish the expression levels of *Dcr-2* in TBPH-mutant backgrounds. For these experiments, transgenic flies containing the *Dcr-2* gene cloned under UAS regulatory sequences (UAS-*Dcr-2*) were crossed against insects carrying the pan-neuronal driver *elav-GAL4* or the more constrained motoneurons promoter *D42-GAL4*. Strikingly, we observed that the expression of UAS-*Dcr-2* in neurons or motoneurons was sufficient to revert the serious locomotive problems showed in of TBPH-null larvae (tbph^Δ23^/tbph^Δ23^; *elav-GAL4* or *D42-GAL4*/UAS-*Dcr-2*) compared to identical flies expressing the unrelated protein GFP (tbph^Δ23^/tbph^Δ23^; *elav-GAL4* or *D42-GAL4*/UAS-GFP) (Figure 4A). Moreover, we found that the recovery of the fly locomotion due to *Dcr-2* expression was followed by the outgrowth of the motoneurons synaptic terminals and the reinnervation of the underlying muscles (Figure 4B and C). These modifications, were followed by the reorganization of the glutamate receptor clusters at the postsynaptic membranes (Figure 4D and E). In addition, we detected that the expression of UAS-*Dcr-2* was able to revert the overexpression of *gypsy* in TBPH-mutant brains (Figure 4F), demonstrating that the alterations in *Dcr-2* levels were responsible for the abnormal activation of *gypsy* and the neurodegeneration associated with defects in TBPH. In addition, our results predict that therapeutic interventions aimed to potentiate *Dcr-2* activity along with the siRNA machinery, would be beneficial to prevent the neurodegeneration occasioned by alterations in TBPH function. In agreement of this idea, we observed that TBPH-minus larvae treated with enoxacin (Shan et al., 2008) were able to recover their locomotive problems and motoneurons synaptic defects revealing that similar therapeutic strategies could be beneficial in patients with ALS (Figure 4G-J).

**Figure 4.**
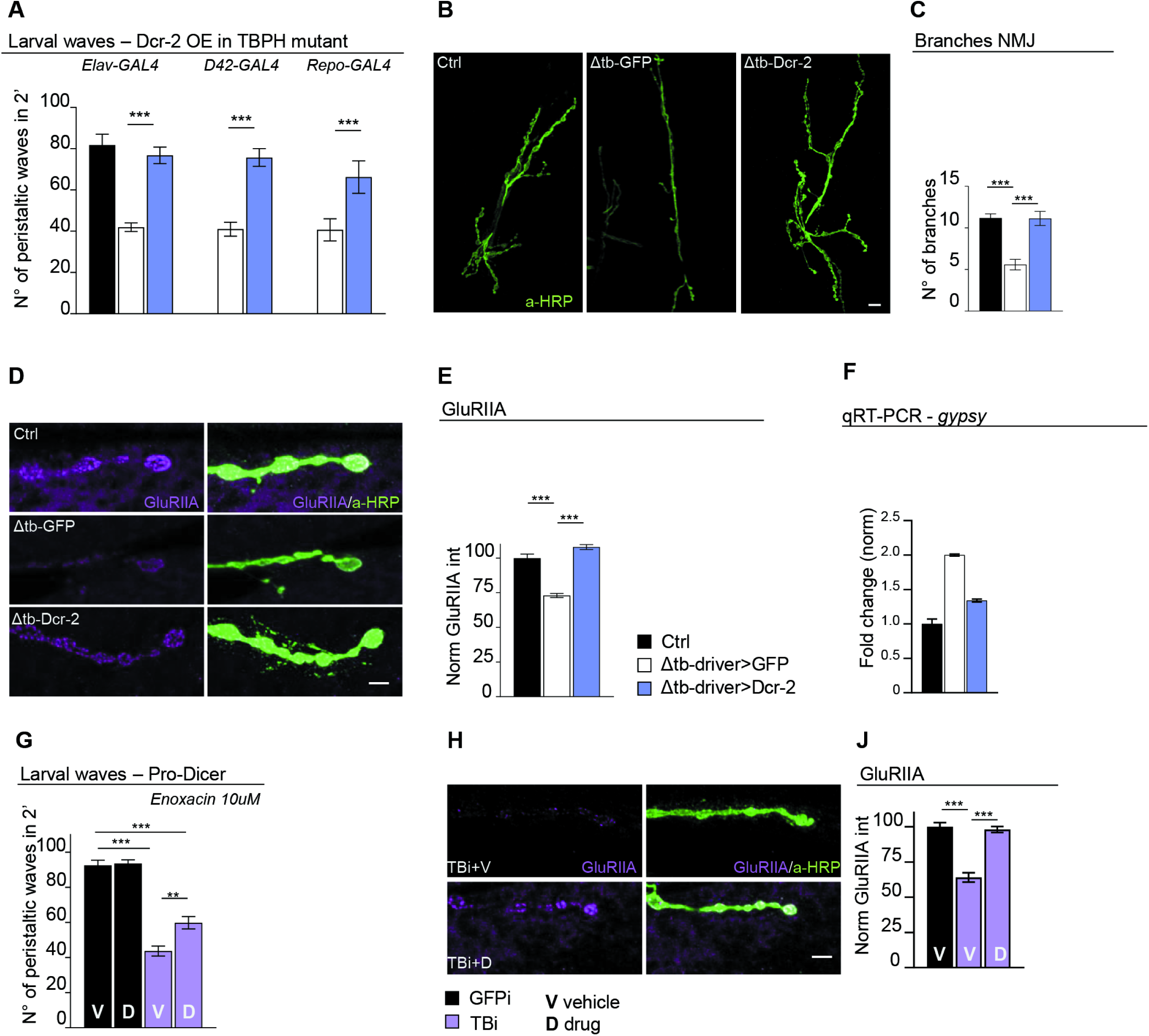
Genetic rescue of *Dcr-2* expression recovers TBPH mutant pathological phenotypes. (**A**) Number of peristaltic waves of Ctrl (*w*^1118^), Δtb-driver>GFP (tbph^Δ23^/tbph^Δ23^;driver-GAL4/UAS-GFP) and Δtb-driver>Dcr-2 (UAS-Dcr-2/+;tbph^Δ23^/tbph^Δ23^;driver-GAL4/+). Elav-GAL4, D42-GAL4 and Repo-GAL4 were used as reported on the figure. *n*=20. (**B**) Confocal images of third instar NMJ terminals in muscle 6/7 second segment stained with anti-HRP (in green) in Ctrl, Δtb-driver>GFP and Δtb-driver>Dcr-2, using elav-GAL4. (**C**) Quantification of branches number. *n*=15. (**D**) Confocal images of third instar NMJ terminals in muscle 6/7 second segment stained with anti-HRP (in green) and anti-GluRIIA (in magenta) in Ctrl, Δtb-driver>GFP and Δtb-driver>Dcr-2, using elav-GAL4. (**E**) Quantification of GluRIIA intensity. *n*>200 boutons. (**F**) Real time PCR of *gypsy* transcript levels normalized on *Rpl11* (housekeeping) in Ctrl, Δtb-driver>GFP and Δtb-driver>Dcr-2, using elav-GAL4. *n*=2, error bars SEM. (**G**) Number of peristaltic waves of GFPi (Dcr-2/+; tbph^Δ23^,elav-GAL4/+;UAS-GFP-IR/+) and TBi (Dcr-2/+; tbph^Δ23^,elav-GAL4/+;UAS-TBPH-IR/+) fed with 10 µM Enoxacin (D) compared to vehicle only (V). *n*=20. (**H**) Confocal images of third instar NMJ terminals in muscle 6/7 second segment stained with anti-HRP (in green) and anti-GluRIIA (in magenta) in GFPi and TBi. (**J**) Quantification of GluRIIA intensity. *n*>200 boutons. **p<0.01, ***p<0.001 calculated by one-way ANOVA, error bars SEM. Scale bar 10µm (in B) and 5µm (in D and H).

## DISCUSSION

The activation of retrotransposons (RTEs) was observed in brain tissues obtained from patients affected from distinctive neurodegenerative diseases and, alterations in the regulation of these elements were described in patients carrying familial or sporadic mutations in TDP-43 suggesting that these events might be related. In agreement with this hypothesis, the overexpression of human TDP-43 provoked the dysregulation of RTEs and neurodegeneration in different animal models (Krug et al., 2017; Li et al., 2013). Yet, these experiments were largely base on the aberrant expression of a neurotoxic protein, making difficult to determine if the results reported were due to a toxic gain function effect of TDP-43 or the dominant interference of this protein with nonspecific mRNA targets and proteins partners. Therefore, remains a matter of debate if the physiological function TDP-43 is required to maintain the repressed status of RTEs or whether the activation of RTEs contribute to the neurodegeneration induced by defects in the function of endogenous TDP-43. In order to find answers to these questions, we performed a genome wide analysis using DNA microchips hybridized with head tissues obtained from null alleles of TBPH, the TDP-43 homolog protein Drosophila. As a result, we found a number of RTEs that appeared consistently dysregulated in tbph^Δ23^ and tbph^Δ142^ mutant flies and these positive hits were further confirmed by quantitative RT-PCR. Interestingly, we observed that one of the most upregulated RTEs was the *gypsy* and detected that the glycoprotein *env*, codified for this retrotransposon, was also upregulated in TBPH-mutants heads (Song et al., 1994). The Drosophila *env* protein, presents strong homology with the human glycoprotein ERV codified by the endogenous retrovirus-K (HERV-K) and, more interestingly, these viral transcripts were found accumulated in the CNS of patients with ALS, suggesting that the regulatory mechanisms behind the activation of these elements might be conserved and present in the physio pathogenesis of ALS. In addition, we detected that the genetic rescue of the missing copies of TBPH was able to repress the RTEs activation as well as the accumulation of the *env* protein in TBPH-null backgrounds demonstrating that these alterations were specific.

### The activation of RTEs provoke motoneurons degeneration in TBPH-null flies

Regarding to the biological implications of the results described above, several lines of investigation have suggested that the mobilization of the RTEs provokes neuronal decline and degeneration (Guo et al., 2018; Krug et al., 2017; Li et al., 2013). On the contrary, parallel studies have reported that the activation of the retrotransposons drives genomic heterogeneity and promotes neurogenesis in flies (Bodea Gabriela O. et al., n.d.). Taking into account these possible scenarios, we found that the suppression of transposons retrotranscription through the administration of revert transcriptase inhibitors and/or nucleoside revert transcriptase inhibitors, was able to ameliorate the locomotive problems described in TBPH-minus flies. More specifically, we observed that the suppression of *gypsy* in neurons or motoneurons was sufficient to revert locomotive defects, promote motoneurons terminals growth and prevent muscles denervation in Drosophila TBPH-null mutants. These results, imply TBPH is physiologically required to prevent the neurotoxic activation of these transposable elements in neurons and, more restrictedly, in motoneurons. On the contrary, the silencing of *gypsy* in glial cells was not able to rescue the phenotypes described in TBPH-null flies suggesting that the repression of the retrotransposon in these tissues is not sufficient to prevent the neurodegeneration induced by the activation of *gypsy*.

### TBPH prevents RTEs-mediated neurodegeneration via the regulation of *Dcr-2* levels

At the molecular level, we found that the RNA silencing activity of the siRNA machinery was reduced in TBPH-null neurons. Additionally, we detected that *Dcr-2*, one of the principal components of the siRNA pathway, was downregulated in TBPH-mutant heads suggesting that defects in the activity of this endoribonuclease might be responsible for the alterations in RTEs repression described above. In agreement with these hypotheses, we observed that the genetic rescue of *Dcr-2* expression was able to prevent the activation of the *gypsy* as well as rescue motoneurons degeneration in TBPH-loss-of-function Drosophila. Furthermore, we established that TBPH forms molecular complexes with *Dcr-2* through physically interactions with the mRNA and the protein itself. The formation of similar protein complexes together with Dicer and Drosha were described for human TDP-43 (Kawahara and Mieda-Sato, 2012), suggesting that these mechanisms might be conserve and present in ALS. In agreement with this idea, we found that the suppression of TDP-43 induce the downregulation of Dicer in human neuroblastoma cell lines indicating that TDP-43 function is required to prevent defects in Dicer activity.

### Pharmacological treatments aimed to enhance *Dcr-2* activity were able to rescue motoneurons degeneration in TBPH-null flies

Finally, our experiments demonstrated that TBPH physically interacts with *Dcr-2* in mRNA and the protein complexes signifying that TBPH may act as a regulatory component of the RNA-induced silencing complexes (RISC) in Drosophila neurons. In consonance with these findings we uncovered that pharmacological treatments utilizing compounds capable to activates the siRNA pathway like enoxacin, were able to restore the locomotive behaviors and the formation of neuromuscular synapsis in TBPH-deficient flies. These therapeutic interventions, either alone or in combination with NRTIs and NNRTIs, may help to control the progress of the disease in patients with familial or sporadic ALS.

## MATERIAL AND METHODS

### Fly strains and maintenance

All flies were maintained at 25°C, with a 12:12 hour light:dark cycle, on standard cornmeal food (agar 6.25 g/L, yeast 62.5g/L, sugar 41.6 g/L, flour 29 g/L, propionic acid 4.1ml/L).

The genotype of the flies used in this work are indicated below:

*w*^1118^ - w;tbph^Δ23^/CyO^GFP^ - w;tbph^Δ142^/CyO^GFP^ - w;elav-GAL4/CyO^GFP^ - w;;D42-GAL4 - Repo-GAL4/TM3,Sb – GMR-GAL4/CyO - UAS-Dcr-2 - w;;UAS-EGFP - w;UAS-TBPH - w;;UAS-TBPH^F/L^ - UAS-gypsy-IR insertion 3 and 4 (gifted by Professor Peng Jin) - UAS-TBPH-RNAi/TM6b (#ID38377, VDRC) - UAS-EGFP/TM3,Sb - UAS-GFP-IR (#9330 Bloomington).

### Larval movement

Peristaltic waves of third instar larvae were performed as already described in (Feiguin et al., 2009). Briefly, larvae, after genotype selection, were rinsed in water and transferred to a 0.7% agarose dish (94mm diameter) and peristaltic waves were counted for a period of two minutes. A minimum of 20 animals was analyzed for each genotype to reach a statistical representative population.

### Drug treatment of larvae

Parental fly crosses were settled on standard cornmeal added of the below listed drugs with the reported final concentration: Stavudine 10μM, Azidotimidine 10μM, Tenofovir 10μM, Abacavir (#SML0089 Sigma) 10μM, Rilpivirine (#10410 Sigma) 10μM, Enoxacin (#AB143281 Abcam) 10μM, Lamivudine (#L1295 Sigma) 10μM. For each drug a vehicle-only control group was arranged. Parental flies have been maintained 24 hours in the tubes to allow the embryo laying. Synchronized embryos were grown to obtain third instar larvae to be tested for mobility or to be analyzed NMJ morphology.

### RNA extraction and microarray analysis

RNA, both from adult dissected brains and adult heads, was extracted with RNeasy Microarray tissue kit (QIAGEN #73304). Gene expression analysis was performed on three independent biological replicates by GenoSplice company on the Affimetrix platform using Gene Chip Drosophila Genome 2.0 Array. RNA extracted from Drosophila adult heads, 1 day aged and sex-matched, of both wild type and TBPH null alleles (*tbph*^Δ23^ and *tbph*^Δ142^) were subjected to quality control tests before chip hybridization. The min-fold change for both up-regulated and down-regulated genes was settled to 1.5.

### Immunohistochemistry, confocal acquisition and quantification

Third instar larvae body were dissected and stained as previously described (Romano et al., 2014). Larvae were dissected in HL-3, fixed in 4% paraformaldehyde 20 minutes (5 min in methanol for anti-GluRIIA) and subsequently blocked in 5% Normal goat serum (Vector laboratories #S-1000) in PBS, 0.1% Tween 20. Primary antibody incubations were performed over night at 4°C, while secondary antibodies were incubated at room temperature for 2 hrs. Dilutions of the antibodies used are reported below: anti-HRP (Jackson 1:150), anti-GluRIIA 8B4D2c (DSHB 1:15), Alexa-Fluor® 488 (mouse or rabbit 1:500) and Alexa-Fluor® 555 (mouse or rabbit 1:500).

Stained larvae were mount with Slow fade Gold (#S36936 Thermo Fisher Scientific) and images of muscles 6 and 7 of the second abdominal segments were gained on a Zeiss LSM880 Laser scanning microscope (63x oil lens). All acquisitions performed in these experiments were simultaneously processed using the same microscope settings and subsequently analyzed by ImageJ (Wayne Rasband, NIH) and Prism (GraphPad, USA) software.

### Cell culture and RNA interference

SH-SY5Y neuroblastoma cell line was cultured in DMEM-Glutamax (#31966-021, Thermo Fisher Scientific) supplemented with 10% fetal bovine serum and 1X antibiotic-antimycotic solution (#A5955; Sigma). For RNA interference 2-4×10^5^ cells were seeded in a 60mm plate in 2ml of medium containing 10% fetal serum. Two rounds of silencing, for a total of 48 hrs silencing, were carried out. HiPerfect Transfection Reagent (#301705, Qiagen) and Opti-MEM I reduced serum medium (#51985-026, Thermo Fisher Scientific) were used with a 200nM final concentration of siRNA, (TDP43: 5’-gcaaagccaagaugagccu-3’ and EGFP control: 5’-gcaccaucuucuucaagga-3’; Sigma). Silenced cells were collected by trypsinization, lysed in RIPA buffer and immunoblotted.

### Immunoblot

Drosophila adult heads or brains were homogenized in lysis buffer 1X (10mM Tris, 150mM NaCl, 5mM EDTA, 5mM EGTA, 10% Glycerol, 50mM NaF, 5mM DTT, 4M Urea, pH 7.4, plus protease inhibitors and protein content quantified with Quant-iT Protein Assay Kit (#Q33211 Thermo Fisher Scientific). SH-SY5Y cells, were homogenized in RIPA buffer (NaCl 150mM, NP-40 1%, Sodium Deoxycholate 0.5%, SDS 0.1%, EDTA 2mM, Tris 50mM, pH8.0) added of protease inhibitors and protein lysates were quantified by Pierce™ BCA Protein Assay Kit (#23225, Thermo Fisher Scientific). Lysates were separated on SDS-PAGE and wet-transferred to nitrocellulose membranes (#NBA083C, Whatman). The primary antibody used were: anti Env (1:100 gifted by Prof. Christophe Terzian), anti Dcr-2 (1:300 #ab4732, ABCAM), anti h-Dicer (1:3000, #PA5-78446 Thermo Fisher Scientific), anti hTDP (1:4000, #12892-1-AP, Proteintech), anti GFP (1:3000, #A11122 Thermo Fisher Scientific), anti-TBPH (1:4000, homemade, (Feiguin et al., 2009), anti-GAPDH (1:1000 #sc-25778, Santa Cruz), anti-Tubulin (1:2000, #CP06, Calbiochem).

### Immunoprecipitation for protein-protein interaction

Approximately one hundred Drosophila heads for each genotype (GMR-GAL4/UAS-TBPH and GMR-GAL4/+) were collected by flash freezing and homogenized in immunoprecipitation buffer (20mM Tris pH7.5, 110mM NaCl, 0.5 % Triton X-100, and protease inhibitors (Roche #11836170001)) with a Dounce homogenizer. Lysates were subjected to 0.4g centrifugation for 5 minutes to remove largest debris and protein content quantified by BCA (#23225, Thermo Fisher Scientific). Equal protein amounts were added of protein G magnetic beads (#10003D, Thermo Fisher Scientific) coated with anti FLAG-M2 antibody (#F3165, Sigma). After an overnight incubation on rototor at 4°C, beads were subjected to washes and finally heated 70°C for 10 minutes in 1XSDS-PAGE loading dye to elute immunoprecipitated proteins that were subsequently immunoblotted with anti-TBPH and anti-Dicer2.

### Immunoprecipitation for RNA enrichment

Drosophila heads collected by flash freezing in liquid nitrogen (elav-GAL4/UAS-TBPH and elav-GAL4/+;UAS-TBPH^F/L^/+) were homogenized in immunoprecipitation buffer (20mM Hepes, 150mM NaCl, 0.5mM EDTA, 10% glycerol,0.1% Triton X-100, and 1mM DTT plus protease inhibitors (Roche #11836170001)) with a Dounce homogenizer and the lysate subjected to 0.4g centrifugation for 5 minutes to remove largest debris. Cleared lysates were added of protein G magnetic beads (#10003D, Thermo Fisher Scientific) coated with anti FLAG-M2 antibody (#F3165, Sigma) and incubated 4°C for half an hour. After five washes with immunoprecipitation buffer, beads were TRIZOL (#15596-026, Ambion) treated to extract RNA.

### qRT-PCR

RNA was DNAse treated with TURBO DNA-*free*™ Kit (#AM1907, Thermo Fisher Scientific) and retrotranscribed with Oligo(dT)_20_ Primer (#18418020, Thermo Fisher Scientific) and Superscript III Reverse Transcriptase (#18080-093, Thermo Fisher Scientific). Real time PCR was performed with Platinum SYBR Green qPCR SuperMIX UDG (#11733-038, Thermo Fisher Scientific) on a Bio-Rad CFX96 qPCR System. Below the used primers:

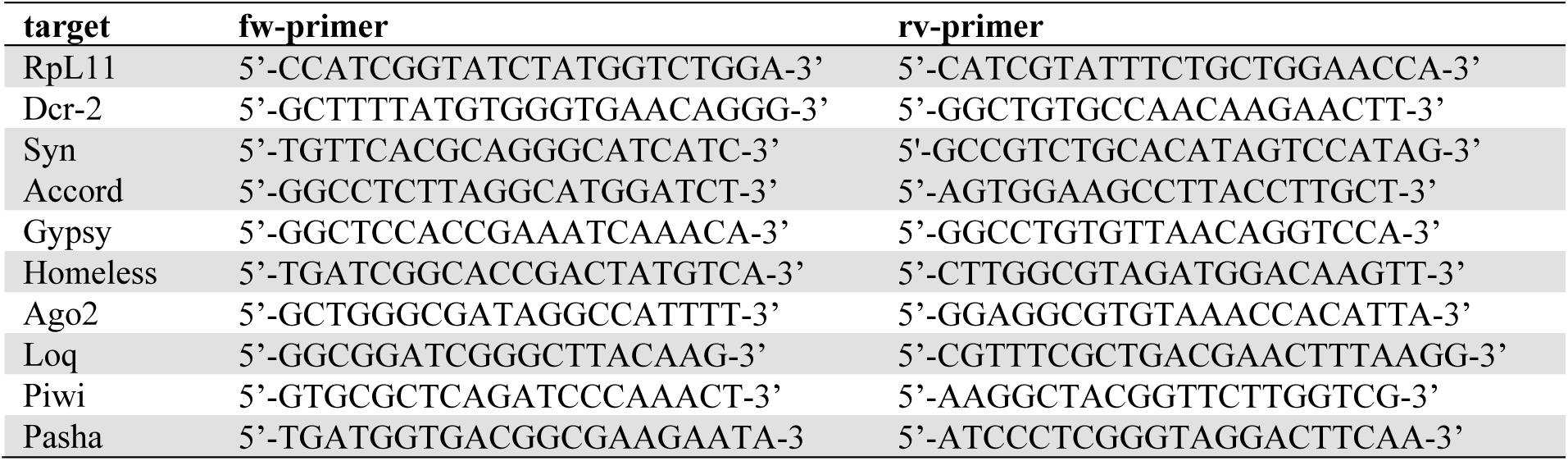

### Statistical analysis

All statistical analysis was performed with Prism (GraphPad, USA) version 6.0. One way Anova with Bonferroni correction was applied as statistical test. In all figures all the values were displayed as the mean and the standard error of the mean (SEM). Statistical significance was displayed as *p<0.05, **p<0.01, ***p<0.001.

## ACKNOWLEDGEMENTS

We thank professor Christophe Terzian, Dr. Franck Touret and Dr. Barbara Viginier to provide us anti-ENV polyclonal antibody and professor Peng Jin to provide us gypsy transgenic flies. The Bloomington Stock Center and Developmental Studies Hybridoma Bank for stocks and reagents.

## Conflict of Interest statement

The authors declare none conflict of interest.

## FUNDING

The present work was supported by ARISLA (CHRONOS) and BENEFICENTIA Stiftung.

## Supplementary Material

**Figure 1-figure supplement 1.**
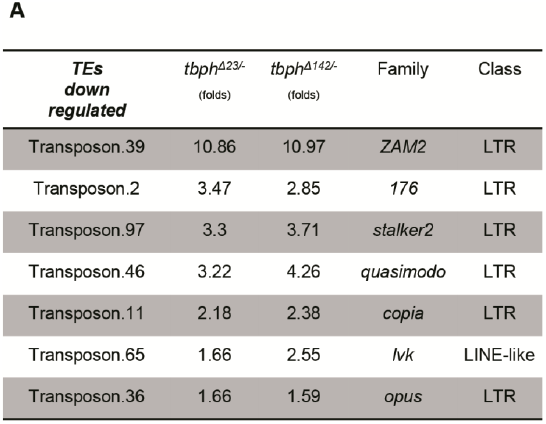
Microarray results of downregulated TEs in TBPH mutants: the fold change of TEs was reported for both tbph mutant alleles (Δ23 and Δ142) referred to *w*^1118^; TEs family and class were also indicated.

**Figure 2-figure supplement 1.**
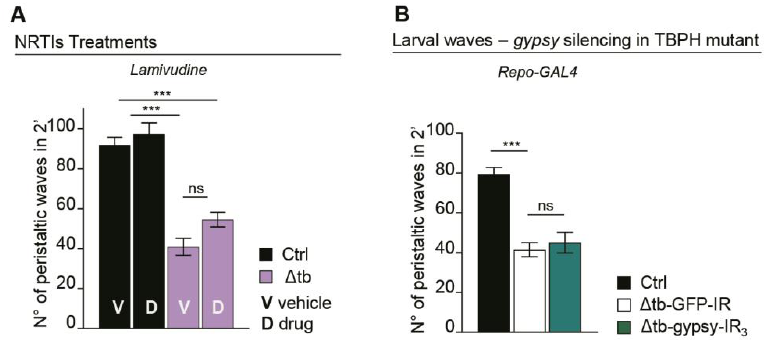
(**A**) Number of peristaltic waves of Ctrl (*w*^1118^) and Δtb (tbph^Δ23^/ tbph^Δ23^) fed with NRTIs drugs (D) compared to vehicle only (V). *n*=20. (**B**) Number of peristaltic waves of Ctrl (*w*^1118^), Δtb-GFP-IR (tbph^Δ23^ /tbph^Δ23^; Repo-GAL4/UAS-GFP-IR) and Δtb-gypsy-IR3 (tbph^Δ23^/tbph^Δ23^; Repo-GAL4/UAS-gypsy-IR3). *n*=20. ns=not significant, ***p<0.001 calculated by one-way ANOVA, error bars SEM.

**Figure 3-figure supplement 1.**
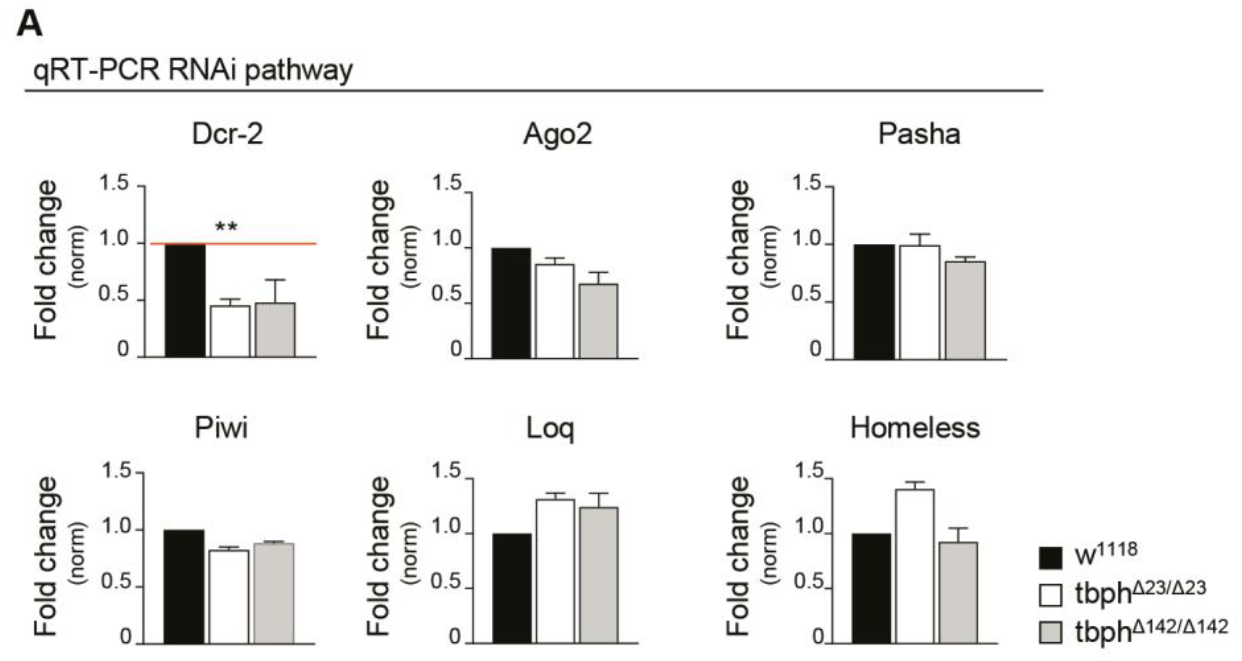
Real time PCR of *Dicer-2* (*Dcr-2), Argonaute 2 (Ago2), Pasha, Piwi, Loquacious (Loq)* and *Homeless* transcript levels normalized on *Rpl11* (housekeeping) in adult heads of *w*^1118^, tbph^Δ23^/tbph^Δ23^ and tbph^Δ142^/tbph^Δ142^. *n*=2, error bars SEM.

